# Free fatty acid receptor 4 agonists stimulate insulin secretion via different mechanisms in mouse versus human islets

**DOI:** 10.1101/2025.08.15.670586

**Authors:** Laura Reininger, Muhammad Rehman, Amélia Bouabcha, Sarah Ferragne, Caroline Tremblay, Mélanie Ethier, Michelle E. Kimple, Julien Ghislain, Mark O. Huising, Vincent Poitout

**Affiliations:** University of Montreal Hospital Research Center (CRCHUM), Montreal, QC, Canada; University of Wisconsin School of Medicine and Public Health, Madison, WI, USA; Department of Neurobiology, Physiology & Behavior, College of Biological Sciences, University of California Davis, Davis, CA, USA; Department of Physiology and Membrane Biology, School of Medicine, University of California Davis, Davis, CA, USA; Department of Medicine, University of Montreal, Montreal, QC, Canada

**Keywords:** FFAR4, GPR120, insulin, somatostatin, Gα_z_, islets

## Abstract

The free fatty acid receptor FFAR4 is expressed in pancreatic islets, and its activation potentiates insulin and inhibits somatostatin (SST) secretion. We investigated the mechanisms of action of FFAR4 on hormone secretion in mouse and human islets. The effects of the FFAR4 agonist Compound A (Cpd A) on insulin and SST secretion were investigated in islets from mice following ablation of δ cells, deletion of SST and deletion of the G protein Gα_z_ (*Gnaz^-/-^*), in purified mouse β and δ cells, in human EndoC-βH5 cells, and in human islets. Ca^++^ dynamics in response to Cpd A were measured in δ cells from *Gnaz^-/-^* mouse islets and in human islets. The insulinotropic effect of Cpd A was lost in δ cell-ablated and SST-deficient mouse islets and was absent in purified mouse β cells. Gα_z_ deletion prevented Cpd A inhibition of SST secretion but not the potentiation of insulin release. Cpd A diminished Ca^++^ transients in mouse δ cells, an effect that was lost in Gα_z_ deficient islets. In human islets, FFAR4 activation increased insulin secretion and intracellular Ca^++^ transient independent of SST secretion. Consistent with a direct effect on β cells, Cpd A potentiated insulin secretion in human EndoC-βH5 cells. We conclude that FFAR4 activation stimulates insulin secretion from mouse islets indirectly via Gα_z_-coupled inhibition of SST secretion from δ cells, while in human islets, it stimulates insulin release via a direct effect on β cells. These key species-related differences are to be taken into account as FFAR4 is considered a potential therapeutic target for metabolic diseases.

**STRUCTURED ABSTRACT:** *Objectives:* The free fatty acid receptor FFAR4 (GPR120) is expressed in the murine islet where its activation promotes insulin and glucagon and inhibits somatostatin (SST) secretion. However, its precise mechanism of action in different islet cells is still unknown, and potential species-related differences have not been explored. This study was aimed to address three questions: 1-What is the relative importance of δ cells and SST in the insulinotropic effect of FFAR4 in mouse islets? 2-Which G protein does FFAR4 couple to in mouse δ cells? 3-Does FFAR4 stimulate insulin secretion by similar mechanisms in mouse and human islets?

*Methods:* The effects of the FFAR4 agonist Compound A (Cpd A) on insulin and SST secretion were investigated in islets from mice following ablation of δ cells, deletion of SST and deletion of the G protein Gα_z_ (*Gnaz^-/-^*), in purified mouse β and δ cells, in human EndoC-βH5 cells, and in human islets. Ca^++^ dynamics in response to Cpd A were measured in δ cells from *Gnaz^-/-^* mouse islets and in human islets using adenovirally-transduced Ca^++^ reporters.

*Results:* The insulinotropic effect of Cpd A was lost in δ cell-ablated and SST-deficient mouse islets and was absent in purified mouse β cells. Gα_z_ deletion diminished Ca^++^ transients in mouse δ cells, prevented Cpd A inhibition of SST secretion but did not inhibit the potentiation of insulin release. In human islets, FFAR4 activation increased insulin secretion and intracellular Ca^++^ transient without affecting SST secretion. Consistent with a direct effect on β cells, Cpd A potentiated insulin secretion in human EndoC-βH5 cells.

*Conclusion:* FFAR4 activation stimulates insulin secretion from mouse islets indirectly via Gα_z_-coupled inhibition of SST secretion from δ cells, while in human islets, it stimulates insulin release via a direct effect on β cells. These key species-related differences are to be taken into account as FFAR4 is considered a potential therapeutic target for metabolic diseases.

## 1. INTRODUCTION

G protein-coupled receptors (GPCRs) are important drug targets due to their role in many physiological processes and extensive involvement in human pathophysiology [1]. GPCRs transduce signals by engaging heterotrimeric G proteins, composed of Gα, Gβ and Gγ subunits, as well as the G protein-independent β-arrestins [2]. There are four major families of Gα subunits, namely Gα_s_, Gα_i/o_, Gα_q/11_ and Gα_12/13_. The long-chain fatty acid receptor FFAR4 (previously referred to as GPR120) is a class A GPCR considered a potential therapeutic target for the treatment of Type 2 diabetes, as agonists of this receptor exert numerous beneficial effects on glucose and energy homeostasis in preclinical models [3]. FFAR4 activation alleviates chronic systemic inflammation and insulin resistance in obese mice [4; 5], inhibits white adipose tissue lipolysis [6], increases adipogenesis [7–9] and brown adipose tissue thermogenesis [10; 11], suppresses neuroinflammation [12] and regulates food intake [13; 14]. In entero-endocrine cells FFAR4 activation stimulates GLP-1 [15–17], GIP [18; 19] and cholecystokinin [20] secretion and inhibits ghrelin [21–23] and somatostatin (SST) [24] secretion. In pancreatic islets FFAR4 activation prevents lipotoxicity and inflammation [25] and apoptosis [26], promotes insulin [27–33], glucagon [27; 31; 34] and pancreatic polypeptide [35] secretion and inhibits SST [31; 32; 36] secretion.

Pancreatic islets are endocrine organs predominantly composed of β, α and δ cells that, respectively, secrete the major glucoregulatory hormones insulin and glucagon, and locally acting SST. Although glucose is the primary insulin secretagogue, diverse molecules modulate glucose-stimulated insulin secretion (GSIS) [37], either directly via β-cell-specific GPCR activation or indirectly via intra-islet paracrine signaling [38]. The importance of local cell-to-cell communication between islet endocrine cell types in the control of insulin levels has been highlighted by its disruption in Type 2 diabetes [38]. Until recently, how islet FFAR4 activation promotes insulin and glucagon secretion was unknown. A possible indirect role was suggested by the demonstration of elevated *Ffar4* expression in SST-secreting δ cells compared to other islet endocrine cells in mouse islets [31; 32; 35; 36; 39-41]. In a previous study using δ-cell specific *Ffar4* knockout mice, we revealed that the insulinotropic and glucagonotropic effects of FFAR4 activation are mediated by inhibitory signaling in δ cells [31]. We showed that FFAR4 activation in mouse islets leads to a reduction in intracellular Ca^++^ signaling and forskolin-induced cyclic AMP levels in δ cells [31]. Our results are consistent with a model whereby FFAR4 signaling inhibits SST secretion, which in turn alleviates the paracrine inhibitory action of SST and results in an increase in insulin and glucagon secretion. In contrast, Wu et al. [27] and McCloskey et al. [30] showed that FFAR4 agonists stimulate insulin secretion in rodent insulin-secreting cell lines, suggesting a direct effect at least in these β cell lines. Therefore, our field has not yet reached consensus regarding the precise site(s) of action of FFAR4 in islets. Our previous study established a role for SST in FFAR4 action in mouse islets by showing that the non-selective SST receptor antagonist cycloSST partially blocked the insulinotropic and glucagonotropic effects of FFAR4 agonists [31]. However, there remains the possibility of additional, direct mechanisms of action on α or β cells. Of note, a direct effect to stimulate insulin or glucagon secretion would imply coupling to a different G protein – in contrast to the inhibitory actions of FFAR4 on δ cells. To this point, FFAR4 signaling exhibits ligand-dependent differential coupling with Gα_i/o_, Gα_s_ and Gα_q_ and β-arrestin in heterologous cells [^42^]. In mice, FFAR4 stimulates pancreatic polypeptide secretion via coupling to Gα_q_ [^35^] and inhibits gastric ghrelin secretion via coupling to Gα_i/o_ [^23^]. Surprisingly, inactivation of inhibitory Gα_i/o_ proteins using pertussis toxin in mouse islets did not prevent FFAR4 inhibition of SST secretion [31], warranting further investigations into its coupling mechanisms. Finally, although *Ffar4* is predominantly expressed in δ cells in mouse islets, RNA-sequencing studies in human islets identified significant *FFAR4* expression in both β and δ cells [41; 43; 44] suggesting potential species-related differences that would be important to identify as FFAR4 is considered a potential therapeutic target for metabolic diseases.

Given that our understanding of the mechanisms of action of FFAR4 in islets is still fragmented, this study aimed to address the following questions: **1**-What is the relative importance of δ cells and SST in the insulinotropic effect of FFAR4 in mouse islets? **2**-Which G protein does FFAR4 couple to in mouse δ cells? **3**-Does FFAR4 stimulate insulin secretion by similar mechanisms in mouse and human islets?

## 2. MATERIALS AND METHODS

### 2.1 Reagents and solutions

RPMI-1640 and FBS were obtained from Life Technologies Inc (Burlington, ON, Canada). Penicillin/streptomycin was from Multicell Wisent Inc (Saint-Jean Baptiste, QC, Canada). Fatty acid (FA)-free BSA was obtained from Equitech bio (Kerrville, TX). Compound A (Cpd A) was from Cayman Chemical (Ann Arbor, MI). AZ13581827 was generously provided by AstraZeneca (Gothenburg, Sweden). Insulin and glucagon radioimmunoassay (RIA) kits were obtained from MilliporeSigma (St Louis, MO). SST RIA kits were obtained from Alpco Diagnotics (Campbell, CA). Poly-D-Lysine was from Life Technologies Inc. (Burlington, ON, Canada). Prodo Islet Media (PIM(S)) and AB serum (PIM(ABS)) were from Prodo Laboratories (Aliso Viejo, CA). EndoC-βH5 cells, coating solution (βCOAT), media (ULTIβ1 and ULTI-ST) and Krebs buffer solution (βKREBS) for EndoC-βH5 cell culture were obtained from Human Cell Design (Toulouse, France). All other reagents and solutions were from MilliporeSigma unless otherwise specified.

### 2.2 Animals

Animals were handled in accordance with the National Institutes of Health guidelines for the care and use of laboratory animals. Mice were housed under controlled temperatures on a 12-h light/dark cycle with unrestricted access to water and standard laboratory chow. B6N.Cg-*Sst^tm2.1(cre)Zjh^*/J (*Sst^+/Cre^*) on a C57Bl/6N background were purchased from Jackson laboratory (Strain #018973, Bar Harbor, ME). Experimental animals (*Sst^+/+^* and *Sst^Cre/Cre^*) were generated from crosses between *Sst^+/Cre^* mice. C57BL/6-*Gt(ROSA)26Sor^tm1(HBEGF)Awai^*/J (*ROSA26^+/iDTR^*) mice on a mixed C57Bl/6JN background were purchased from Jackson laboratory (Strain #007900) and backcrossed for 9 generations to C57Bl/6N mice. Experimental animals (*Sst^+/Cre^*;*ROSA26 ^+/iDTR^*) were generated from crosses between *Sst^+/Cre^* and *ROSA26 ^iDTR/iDTR^* mice. Diphtheria toxin (DT) (126 ng/200 μl 0.9% saline, List Biological Laboratories, Catalog # 150) or vehicle (0.9% saline) was injected intraperitoneally on days 0, 3, and 4. On day 5 pancreata were processed for immunochemistry or islets isolated for secretion assays. *Gnaz^tm1Iah^*(*Gnaz^+/-^*) on a C57Bl/6N background were kindly provided by Michelle Kimple [45]. Experimental animals (*Gnaz^+/+^*, *Gnaz^+/-^* and *Gnaz^-/-^*) were generated from crosses between *Gnaz^+/-^* mice. Genotyping primers are listed in **Supplementary Table 1**. Animals were born at the expected Mendelian ratios. C57Bl/6N male mice were purchased from Charles River Laboratories (Saint-Constant, QC, Canada).

### 2.3 Islet isolation and cell culture

Mouse islets were isolated at 10-12 weeks of age by collagenase digestion and dextran density gradient centrifugation as described [46] and allowed to recover overnight in RPMI 1640 supplemented with 10% (v/v) FBS, penicillin/streptomycin (100 U/ml), and glucose (11 mM). Upon reception, human islets were allowed to recover overnight in PIM(S) islet media containing 5% (v/v) PIM(ABS) serum, 1% (v/v) glutamine/glutathione mixture and penicillin/streptomycin (100 U/ml).

EndoC-βH5 cells were cultured as described [47]. Frozen cells were thawed in a 37 °C water bath, washed with ULTIβ1 media and seeded on βCOAT coated 24-well plastic plates (175,000 cells/well) and cultured in ULTIβ1 media. On day 6 the media was replaced with ULTI-ST media for 24 h prior to experimentation.

### 2.4 Flow cytometric sorting of mouse β and δ cells

Flow cytometric sorting of mouse islet β and δ cells based on expression of the cell surface markers CD24, CD49f, CD71 and CD81 was performed as described [48] with minor modifications. Batches of 400 islets were dispersed by gentle pipetting for 10 min at 37 °C in Accutase (500 μl, Innovative Cell Technologies, Inc., San Diego, CA). Dispersed islet cells were incubated with antibodies diluted in HBSS supplemented with 2% FBS (Staining medium) for 15 min at 4 °C. Antibodies and dilutions are indicated in **Supplementary Table 2**. Prior to flow cytometric sorting, cells were labeled with LIVE/DEAD Fixable Aqua vivid (405 nm) Dead Cell Stain (BD Biosciences, San Jose, CA) diluted in Staining medium. Cell-sorting was performed using a FACSAria III (BD Biosciences) and data analyzed using FlowJo 9.9.4 (RRID: SCR_008520). Cells sorted for qPCR were collected directly in RLT buffer (Qiagen, Hilden, Germany).

### 2.5 Static incubations for hormone secretion

#### Isolated islets

Mouse and human islets were incubated in KRBH (pH 7.4) supplemented with 0.1% (w/v) FA-free BSA (KRBH/BSA) and 2.8 mM glucose for 20 min. Then, triplicate batches of 20 islets each were dispersed into a 24-well plate and incubated for an additional 20 min in KRBH/BSA and 2.8 mM glucose, followed by a 1-h static incubation in KRBH/BSA in the presence of 2.8 or 16.7 mM glucose and the FFAR4 agonists Cpd A (20 μM) and AZ13581827 (10 μM) or vehicle (EtOH), as indicated in the figure legends. The synthetic FFAR4 agonists Cpd A [5] and AZ13581827 [16] were chosen among the different FFAR4 agonists for their selectivity towards FFAR4.

#### Sorted β and δ cells

Sorted β (200,000/well) and δ (1000/well) cells were suspended in RPMI 1640 supplemented with 10% (wt/vol) FBS, penicillin/streptomycin (100 U/ml), and glucose (11 mM) and plated in poly-D-lysine coated 24-well plates and recovered overnight. For insulin secretion, each well of sorted β cells was preincubated with KRBH/BSA and 2.8 mM glucose for 1 h followed by a 1h-static incubation in the presence of 2.8 or 16.7 mM glucose and Cpd A (20 μM) or vehicle (EtOH). For SST secretion, each well of sorted δ cells was preincubated with KRBH/BSA and 2.8 mM glucose for 1 h followed by 1h-sequential static incubations in the presence of 2.8 mM glucose, 16.7 mM glucose and finally 16.7 mM glucose and Cpd A (20 μM).

#### EndoC-βH5 cells

EndoC-βH5 insulin secretion assays were performed as described [47] with minor modifications. Cells were preincubated in βKREBS with 0.1% (w/v) FA-free BSA (βKREBS/BSA) and 2.8 mM glucose for 1 h. Then, sequential 40 min-static incubations in βKREBS/BSA and 2.8 mM glucose followed by 16.7 mM glucose and Cpd A (20 μM) or AZ13581827 (10 μM), as indicated in the figure legends, were performed.

#### Hormone measurements

Secreted insulin and SST were measured in the supernatant by RIA. Intracellular insulin content was measured after acid-alcohol extraction.

### 2.6 Perifusions

Human islet perifusions were performed as described [49] with minor modifications. Batches of 100 islets were placed in chambers and perifused in KRBH/BSA containing sequentially 2.8 mM glucose for 20 min, 16.7 mM glucose with Cpd A (20 μM), AZ13581827 (10 μM) or vehicle (EtOH) for 40 min, followed by 2.8 mM glucose for 20 min and finally KCl (30 mM) for 20 min. Samples were collected at 2 min intervals and secreted insulin and SST measured by RIA. Intracellular insulin content was measured at the end of the perifusion after acid-alcohol extraction.

### 2.7 RNA extraction and quantitative RT-PCR

Total RNA was extracted from 200 islets, 60,000 sorted β and 1000 sorted δ cells, using RNeasy micro kits (Qiagen). RNA was quantified by spectrophotometry using a NanoDrop 2000 (Life Technologies) and 50 ng of RNA was reverse transcribed. Real-time PCR was performed using a QuantiTect SYBR Green PCR kit (Qiagen). Data were normalized to cyclophilin A (Ppia) mRNA levels. qPCR primer sequences are listed in **Supplementary Table 3**.

### 2.8 Immunohistochemistry of pancreatic sections

Pancreata were isolated, fixed for 4 hours in 4% paraformaldehyde (PFA) and cryoprotected overnight in 30% sucrose. The tissues were embedded in OCT compound, frozen, sectioned at 8 μm, and mounted on Superfrost Plus slides (Thermo Fisher Scientific, Waltham MA). Antigen retrieval was performed using sodium citrate buffer (pH=ƒ6). Sections were stained for NKX6.1 and SST and nuclei were stained with Hoechst 33342 (Thermo Fisher Scientific). Primary and secondary antibodies and dilutions are listed in **Supplementary Table 4**. Slides were mounted using Vectashield mounting medium (Vector laboratories Inc, Newark CA) and images acquired using a Zeiss AX10 fluorescence microscope (Oberkochen, Germany).

### 2.9 Ca^++^ imaging in mouse and human islets

Recombinant adenoviruses for live Ca^++^ imaging Ad5-CMV-jRGECO1b and Ad5-CMV-GcaMP6s-P2A-NLS-mKate2 were generated using the AdEasy protocol [50]. Isolated *Gnaz^+/+^* and *Gnaz^-/-^*mouse islets and human islets (**Supplementary Table 5**) were transduced with Ad5-CMV-jRGECO1b and Ad5-CMV-GcaMP6s-P2A-NLS-mKate2, respectively, at approximately 100 multiplicity of infection. For imaging, microfluidics chambers were bonded to 35-mm dishes with a glass bottom (MatTek, Ashland, MA). Islets were allowed to adhere to the glass during overnight culture. Continuous perfusion of KRBH at a rate of 200 µl min^−1^ was maintained using the Elveflow microfluidics system, with different treatments adjusted using the Mux distributor. Ca^++^ responses were imaged using a Nikon Eclipse Ti2 using a ×60 lens with oil. Regions of interest (ROI) were drawn around individual mouse δ cells or whole human islets on NIS-Elements v.5.02.01. Fluorescence intensity in each ROI was measured to determine sensor activity over time.

### 2.10 Statistical analyses

Data are expressed as mean ± SEM. Significance was tested using standard one-way ANOVA, Brown-Forsythe and Welch ANOVA and corrections, in cases of variance heterogeneity, or two-way ANOVA with post hoc adjustment for multiple comparisons, as appropriate, using GraphPad Instat (GraphPad Software, San Diego, CA). Tukey or Dunnett post hoc tests were performed as indicated in figure legends. p < 0.05 was considered significant.

### 2.11 Study approval

All procedures involving animals were approved by the Institutional Committee for the Protection of Animal at the Centre Hospitalier de l’Université de Montréal. Islets from nondiabetic human donors were obtained from the National Institute of Diabetes and Digestive and Kidney Diseases–sponsored Integrated Islet Distribution Program (RRID:SCR_014387) at City of Hope (Duarte, CA), NIH grant 2UC4DK098085; the Alberta Diabetes Institute Islet Core at the University of Alberta (Edmonton, AB, Canada) and Prodo Laboratories Inc. (Aliso Viejo, CA). The use of human islets was approved by the Institutional Ethics Committee of the Centre Hospitalier de l’Université de Montréal (protocol # MP-02-2019-7880).

## 3. RESULTS

### 3.1 Delta-cell ablation prevents the potentiation of insulin secretion by Cpd A in male mouse islets

Previously, by knocking out *Ffar4* in SST-expressing cells we demonstrated that FFAR4 signaling in δ cells is necessary for the insulinotropic effects of Cpd A in isolated mouse islets [31]. As FFAR4 signaling in δ cells inhibits SST secretion [31; 36] and SST is a paracrine inhibitor of insulin secretion [51], our results were consistent with the possibility that FFAR4 agonism alleviates SST inhibition of insulin secretion. Yet, SST receptor blockade only partially reduced FFAR4 agonist-dependent potentiation of insulin secretion [31], leaving open the possibility of a role of FFAR4-dependent δ cell-derived signals other than SST. To test whether ablation of δ cells prevents the potentiation of insulin secretion by Cpd A, δ-cell deficient male mice were generated by injecting DT into *Sst^+/Cre^*;*ROSA26 ^+/iDTR^* mice which express the DT receptor (DTR) in SST-expressing cells [52]. DT injection of *Sst^+/Cre^*;*ROSA26^+/iDTR^* mice was previously shown to specifically deplete islet δ cells up to 3 months post-injection with no impact on α or β cell numbers [52]. We confirmed the near-complete absence of SST staining in pancreatic sections of DT injected-*Sst^+/Cre^*;*ROSA26 ^+/iDTR^* mice (**Supplementary Fig. S1**). In 1-h static incubations, high glucose concentrations (16.7 mM) significantly increased insulin secretion in islets from both saline- and DT-injected *Sst^+/Cre^*;*ROSA26 ^+/iDTR^*mice compared to the low (2.8 mM) glucose condition (**Fig. 1A**). Glucose increased SST secretion in islets from saline-injected *Sst^+/Cre^*;*ROSA26 ^+/iDTR^* mice (**Fig. 1B**). As expected, secreted SST levels were near the assay limit of detection in DT-injected *Sst^+/Cre^*;*ROSA26 ^+/iDTR^* mice (**Fig. 1B**). As shown previously [31], the FFAR4 agonist Cpd A (20 μM) significantly potentiated GSIS and inhibited glucose-stimulated SST secretion (GSSS) in control islets (**Fig. 1A & B**). In contrast, the effect of Cpd A on GSIS was absent following δ-cell ablation (**Fig. 1A**). As shown previously [52], insulin content was not affected by δ-cell ablation (1,484 ± 157 ng/ml versus 1,506 ± 135 ng/ml for control; n=5-6; NS).

**Figure 1.**
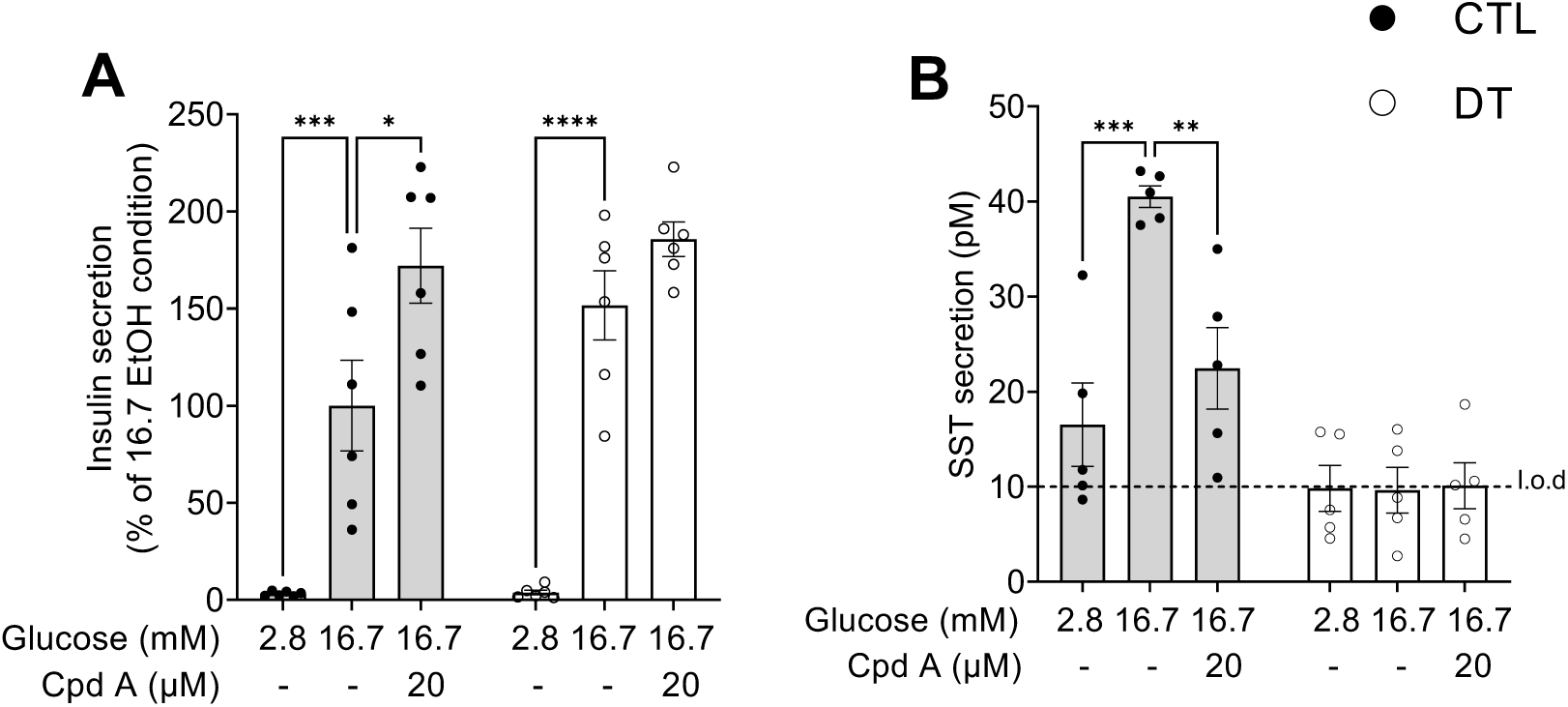
Delta-cell ablation abolishes the potentiation of GSIS in response to Cpd A in male mouse islets. Insulin secretion presented as a percentage of the control 16.7 EtOH condition (A) and SST secretion (B) measured in parallel in 1-h static incubations in response to 2.8 or 16.7 mM glucose with Cpd A (20 μM) or vehicle (EtOH) in isolated islets from *Sst^+/Cre^*;*ROSA26 ^+/iDTR^* male mice injected with saline (CTL) or diphtheria toxin (DT). Data are expressed as mean ± SEM from 5-6 independent experiments. ∗p < 0.05, ∗∗p < 0.005, ∗∗∗p < 0.0005, ∗∗∗∗p < 0.0001 vs 16.7-EtOH condition following two-way ANOVA with Tukey’s post hoc adjustment for multiple comparisons. L.o.d, limit of detection.

### 3.2 Loss of SST expression prevents the potentiation of insulin secretion by Cpd A in female mouse islets

To then specifically ascertain the role of SST in the action of Cpd A in islets, we took advantage of the absence of SST expression in islets from *Sst^Cre/Cre^* mice [52; 53]. We confirmed that homozygous *Sst^Cre/Cre^*mice have no detectable SST protein expression in islets (**Supplementary Fig. S2**). In 1-h static incubations, high glucose concentrations (16.7 mM) increased insulin secretion in islets from both *Sst^+/+^* and *Sst^Cre/Cre^* male (**Fig. 2A**) and female (**Fig. 2B**) mice compared to the control 2.8 mM glucose condition, although statistical significance was only reached in female islets. Glucose also increased SST secretion in islets from *Sst^+/+^*male (**Fig. 2C**) and female (**Fig. 2D**) mice. As expected, SST levels were near the assay limit of detection in *Sst^Cre/Cre^* islets (**Fig. 2C & D**). Surprisingly, in *Sst^+/+^* male mouse islets Cpd A did not significantly potentiate GSIS (**Fig. 2A**), although GSSS was inhibited (**Fig. 2C**). However, Cpd A significantly potentiated GSIS (**Fig. 2B**) and inhibited GSSS (**Fig. 2D**) in control female islets. Importantly, Cpd A had no effect on GSIS (**Fig. 2A & B**) in SST deficient islets. Insulin content was not affected by the constitutive loss of SST in either male (728 ± 108 ng/ml in *Sst^Cre/Cre^*versus 774 ± 79 ng/ml in *Sst^+/+^*; n = 9; NS) or female (1,027 ± 109 ng/ml in *Sst^Cre/Cre^* versus 1,249 ± 137 ng/ml in *Sst^+/+^*; n = 6; NS) islets.

**Figure 2.**
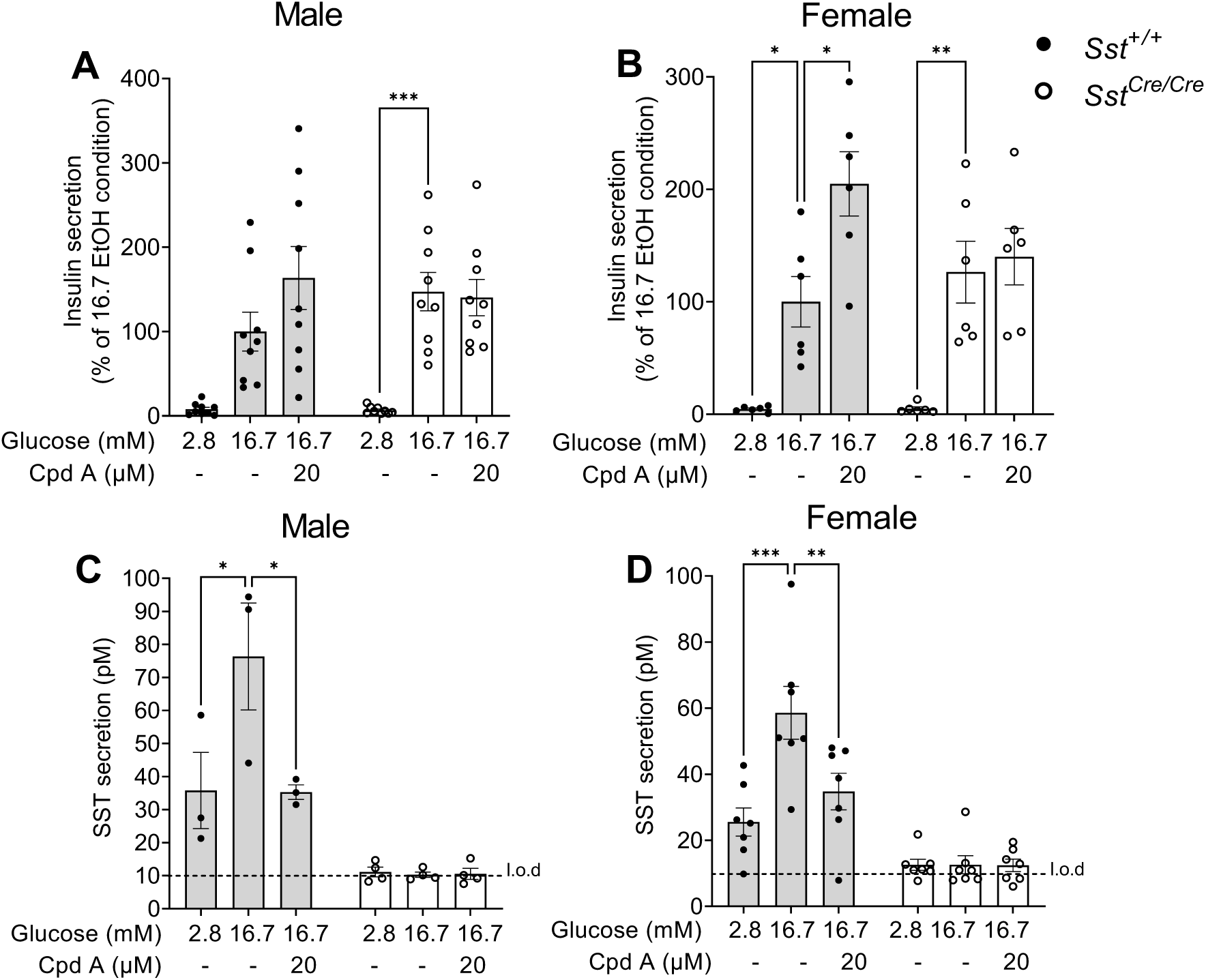
*Sst* inactivation prevents potentiation of GSIS in response to Cpd A in female mouse islets. Insulin secretion presented as a percentage of the control 16.7 EtOH condition (A, B) and SST secretion (C, D) measured in parallel in 1-h static incubations in response to 2.8 or 16.7 mM glucose with Cpd A (20 μM) or vehicle (EtOH) in isolated male (A, C) and female (B, D) islets from *Sst^+/+^* and *Sst^Cre/Cre^* mice. Data are expressed as mean ± SEM from 3-8 independent experiments. ∗p < 0.05, ∗∗p < 0.005, ∗∗∗p < 0.0005 vs 16.7-EtOH condition following two-way ANOVA with Tukey’s post hoc adjustment for multiple comparisons. L.o.d, limit of detection.

### 3.3 Cpd A has no effect on insulin secretion in purified mouse β cells

The data presented above suggest that the increase in insulin secretion in response to FFAR4 agonism in mouse islets is predominantly due to inhibition of SST secretion from δ cells. To confirm this possibility, we purified live α, β and δ cells from wild-type male mouse islets using a sorting strategy described by Berthault et al. [48] (**Supplementary Fig. S3**). Compared to the level of expression of *Ins1*, *Gcg* and *Sst* in whole male mouse islets, flow cytometry-sorted CD24^low^CD81^-^ were enriched in *Ins1* and deficient in *Gcg* and *Sst* mRNA (**Fig. 3A-C**). In contrast, CD24^high^CD49f^-^ were deficient in *Ins1* and *Gcg* and enriched in *Sst* mRNA (**Fig. 3A-C**). CD24^low^CD81^-^ and CD24^high^CD49f^-^ cells are hereafter referred to as sorted β and δ cells, respectively. Insulin and SST secretion were assessed in 1-h static incubations of sorted β and δ cells, respectively, side-by-side with intact islets (**Fig. 3D-G**). As expected, Cpd A significantly potentiated GSIS in intact islets (**Fig. 3D**). In contrast, sorted β cells were unresponsive to Cpd A (**Fig. 3E**). In line with the direct effect of glucose on δ cells [54], glucose increased SST secretion in both intact islets and sorted δ cells (**Fig. 3F & G**). Importantly, Cpd A repressed GSSS in both intact islets and sorted δ cells (**Fig. 3F & G**).

**Figure 3.**
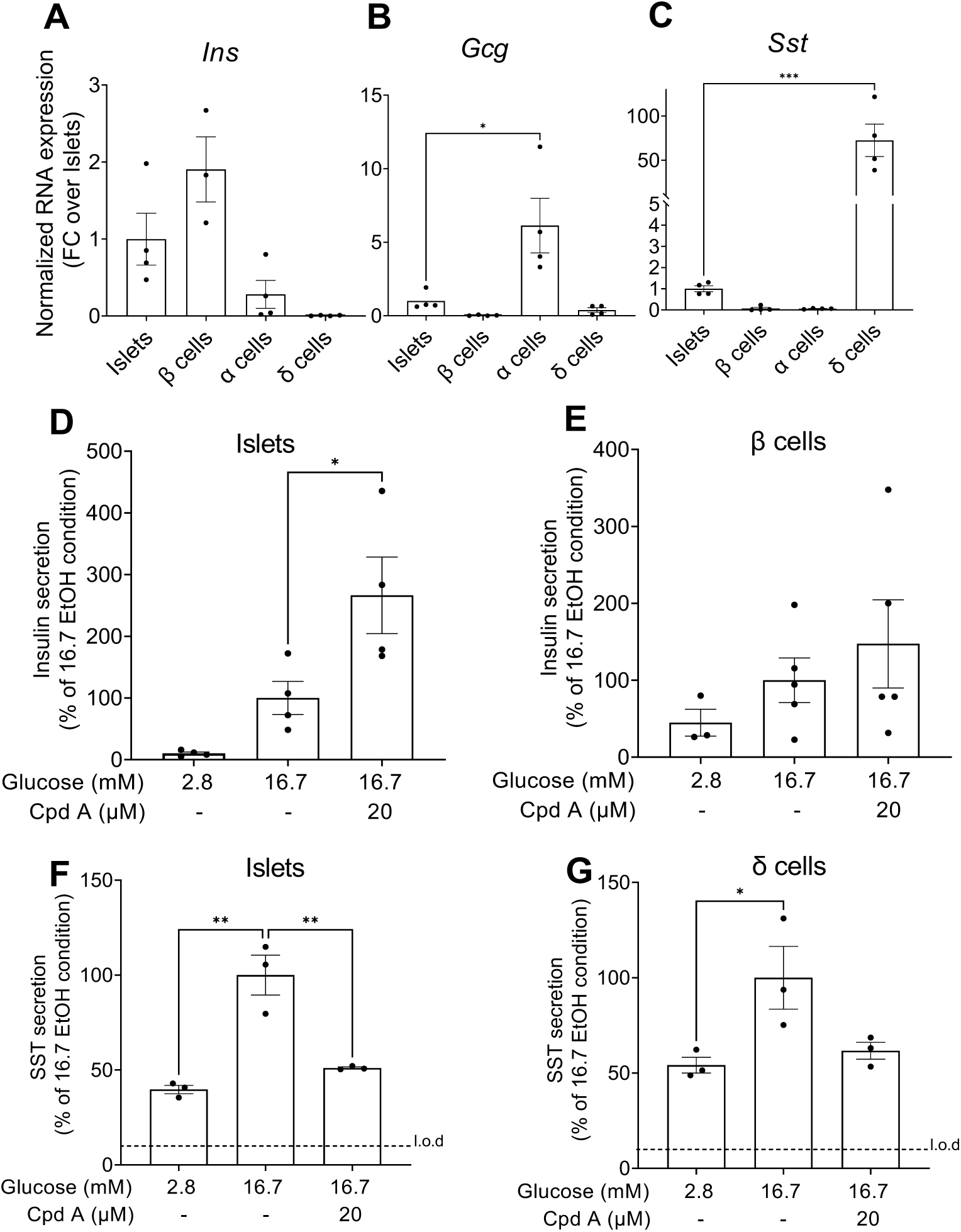
Cpd A does not stimulate GSIS in sorted male mouse β cells. *Ins1* (A), *Gcg* (B) and *Sst* (C) mRNA levels in intact male mouse islets and flow cytometry-sorted CD24^low^CD81^-^ (β), CD24^low^CD81^+^/CD49f^-^ (α) and CD24^high^CD49f^-^ (δ) cells were measured by qPCR and normalized to cyclophilin. Data are presented as the fold change (FC) over intact islets. Insulin (D, E) and SST (F, G) secretion presented as a percentage of 16.7 EtOH condition was assessed in 1-h static incubations in response to 2.8 or 16.7 mM glucose with Cpd A (20 μM) or vehicle (EtOH) in isolated islets (D, F) and sorted β (E) and δ (G) cells from male mice. Data are expressed as mean ± SEM from 3-5 independent experiments. ∗p < 0.05, ∗∗p < 0.005, ∗∗∗p < 0.0005 vs intact islets (A-C) or 16.7-EtOH condition (D-G) using one-way ANOVA with Tukey’s post hoc adjustment for multiple comparisons. L.o.d, limit of detection.

Taken together with our previous studies [31], these data demonstrate that the potentiation of insulin secretion by Cpd A in mouse islets is indirect and mediated by a reduction in SST secretion due to FFAR4-inhibitory signaling in δ cells.

### 3.4 Cpd A inhibition of SST secretion requires the inhibitory G protein Gα_z_ in male mouse islets

FFAR4 was shown to couple to the G proteins Gα_s_, Gα_q_ and Gα_i/o_ in a ligand specific manner in heterologous cells [42]. Since FFAR4 agonism in islets inhibits SST secretion, in our previous study we investigated the involvement of the inhibitory G proteins Gα_i/o_. However, blocking Gα_i/o_ signaling with pertussis toxin did not prevent Cpd A-mediated inhibition of GSSS [^31^]. As Gα_z_ mediates inhibitory signaling in islets and is insensitive to pertussis toxin [^55^], we investigated FFAR4-Gα_z_ coupling in 1-h static incubations of male *Gnaz^+/+^*, *Gnaz^+/-^*and *Gnaz^-/-^* mouse islets [45]. High glucose concentrations (16.7 mM) increased SST secretion in all genotypes, albeit not significantly in *Gnaz^-/-^* islets compared to the control 2.8 mM glucose condition (**Fig. 4A**). In contrast to *Gnaz^+/+^* and *Gnaz^+/-^*islets, Cpd A did not suppress GSSS in *Gnaz^-/-^* islets (**Fig. 4A**). Glucose increased insulin secretion in all genotypes, and Cpd A potentiated GSIS in *Gnaz^+/+^* and *Gnaz^+/-^* islets, as expected. Surprisingly, despite the absence of Cpd A-mediated repression of SST secretion, GSIS was still potentiated by Cpd A in *Gnaz^-/-^* islets (**Fig. 4B**). Islet insulin content was not different between the genotypes (*Gnaz^+/+^*, 998 ± 95 ng/ml; *Gnaz^+/-^*, 1,011 ± 70 ng/ml; *Gnaz^-/-^*, 1,112 ± 84 ng/ml; n = 7; NS).

**Figure 4.**
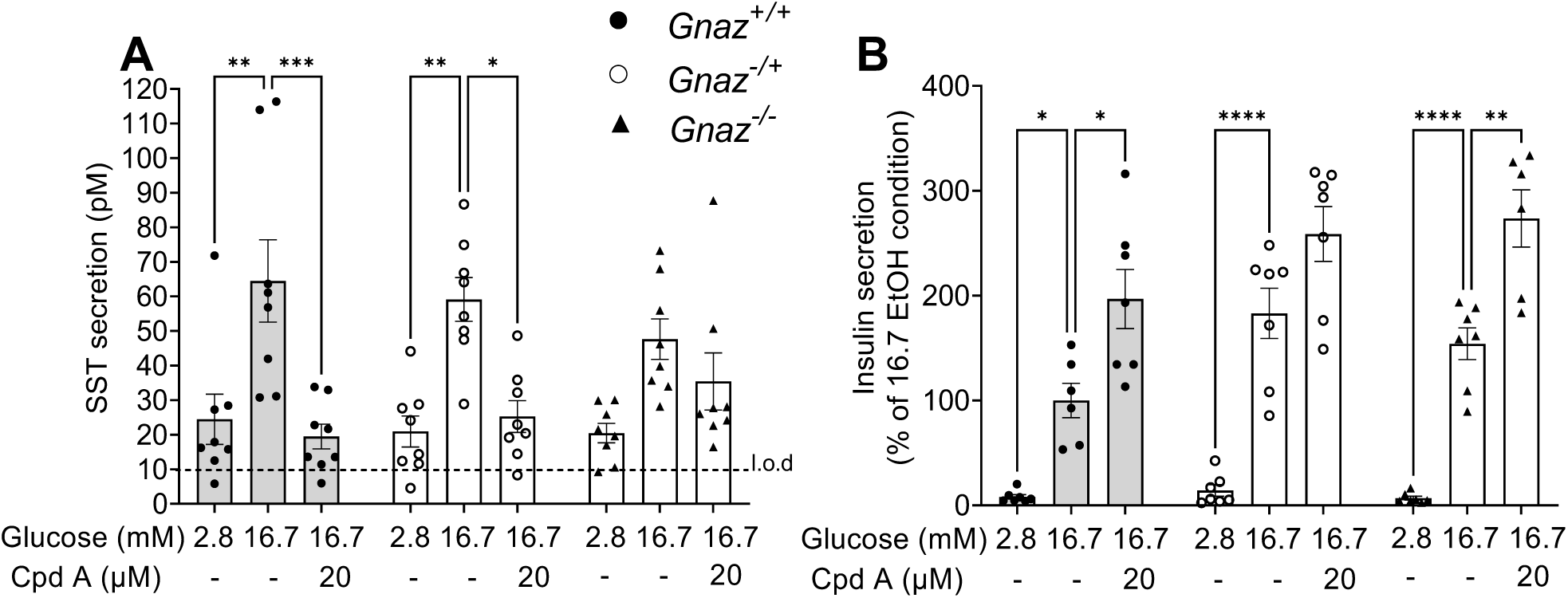
*Gnaz* deletion prevents the inhibition of SST secretion by Cpd A in male mouse islets. SST secretion (A) and insulin secretion measured in parallel and presented as a percentage of the control 16.7-EtOH condition (B) was assessed in 1-h static incubations in response to 2.8 or 16.7 mM glucose with Cpd A (20 μM) or vehicle (EtOH) in isolated islets from *Gnaz^+/+^*, *Gnaz^+/-^*and *Gnaz^-/-^* male mice. Data are expressed as mean ± SEM from 6-7 independent experiments. ∗p < 0.05, ∗∗p < 0.005, ∗∗∗p < 0.0005, ∗∗∗∗p < 0.0001 vs 16.7-EtOH condition following two-way ANOVA with Tukey’s post hoc adjustment for multiple comparisons. L.o.d, limit of detection.

We previously reported that Cpd A decreases δ-cell Ca^++^ signaling in mouse islets [31]. To assess the role of Gα_z_, *Gnaz^+/+^*and *Gnaz^-/-^* mouse islets were infected with an adenovirus expressing the Ca^++^ reporter jRGECO1b. As jRGECO1b was expressed in all islet cells, δ cells were identified by exposure to ghrelin, which stimulates SST secretion [39]. Cpd A diminished Ca^++^ transients in *Gnaz^+/+^* δ cells at 5.5 mM glucose, as expected (**Fig. 5, Supplementary Fig. S4**). In contrast, homozygous deletion of *Gnaz* alleviated the inhibitory effect of Cpd A on δ cell Ca^++^ transients (**Fig. 5, Supplementary Fig. S4**). Ghrelin increased Ca^++^ in both genotypes, albeit with a weaker effect in *Gnaz^-/-^*, confirming the identity of the cells as δ cells.

**Figure 5.**
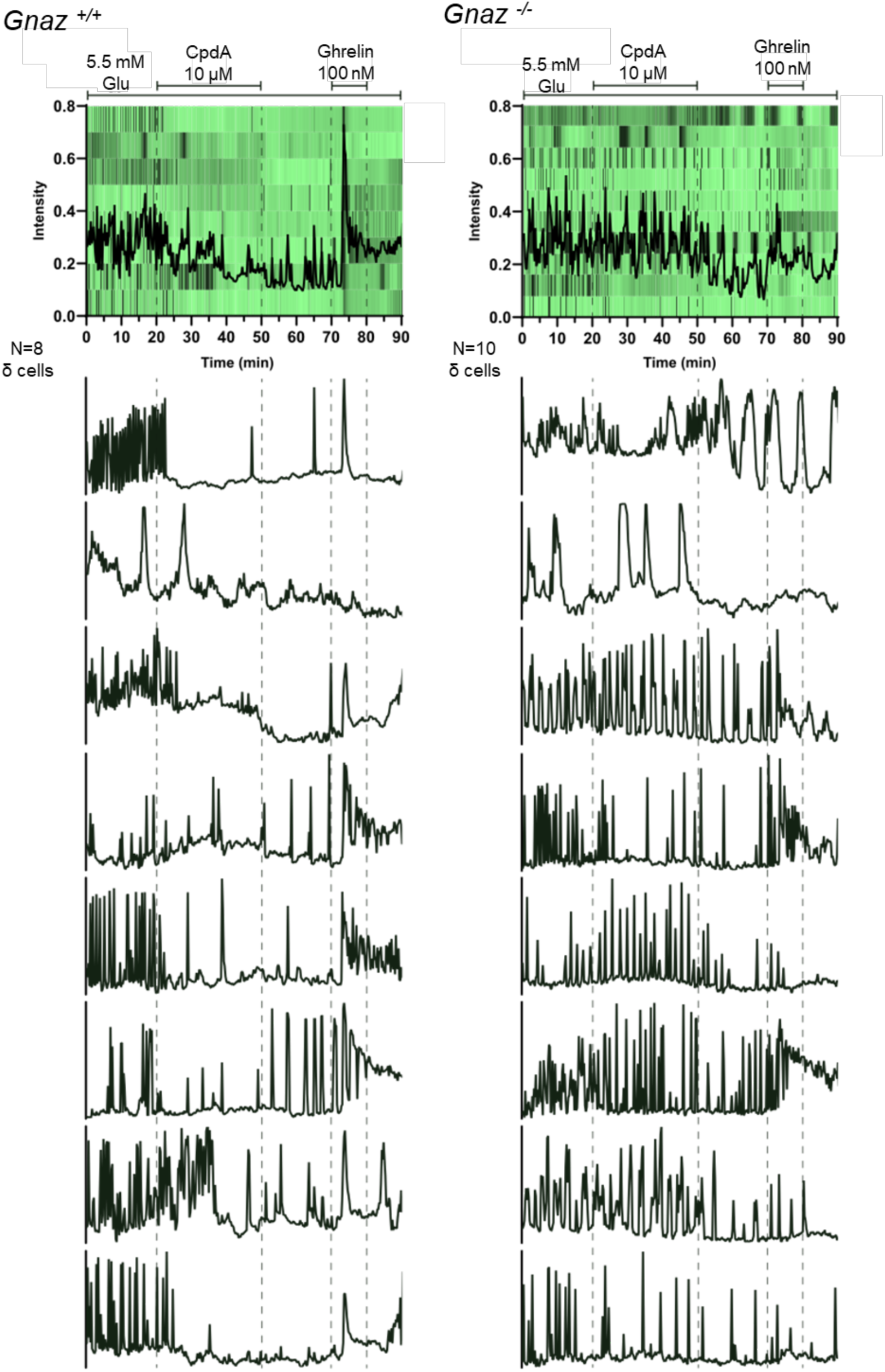
*Gnaz* deletion abrogates the Cpd A-mediated decrease in intracellular Ca^++^ transients in male mouse δ cells. The Ca^++^ sensor jRGECO1b was used to measure Ca^++^ transients in individual δ cells in the presence of 5.5 mM glucose in *Gnaz^+/+^*(left) and *Gnaz^-/-^* (right) male mouse islets. Cpd A (10 μM) was added at the times indicated. Ghrelin (100 nM)-induced Ca^++^ signaling served as a positive control to identify δ cells. Aggregate data of all δ cells analyzed plotted with the average intensity (upper panels) and traces of individual δ cells (lower panels) are shown. Data represent the results of the analysis of 8-10 δ cells in each of one batch of *Gnaz^+/+^*and *Gnaz^-/-^* islets.

Taken together, these data indicate that FFAR4 couples predominantly to Gα_z_ in δ cells to diminish Ca^++^ transients and SST secretion in response to Cpd A, but suggest that Gα_z_ may not be necessary for Cpd A-mediated potentiation of GSIS.

### 3.5 FFAR4 agonists regulate insulin but not SST secretion in human islets

To investigate the role of FFAR4 in human islets we first assessed islet cell-type-specific *FFAR4* expression from single-cell RNA sequencing datasets [44]. Interestingly, whereas ƒ*Ffar4* is predominantly expressed in δ cells in mouse islets [31], significant *FFAR4* expression was also detected in human β cells (**Fig. 6A & B**). We then assessed insulin and SST secretion in adult male and female human islets in response to FFAR4 agonists in 1-h static incubations. Both Cpd A and the structurally dissimilar, FFAR4 specific agonist AZ13581837 potentiated GSIS (**Fig. 6C**). In contrast to mouse islets, SST secretion in response to glucose was not inhibited by either Cpd A or AZ13581837 (**Fig. 6D**). In perifusion experiments, Cpd A provoked a large increase in GSIS in both male and female human islets (**Fig. 6E & F**). GSIS also trended higher in response to AZ13581837, although the area under the curve was not significantly different (**Fig. 6E & G**). Interestingly, following washout of the FFAR4 agonists, KCl potentiated insulin secretion in islets treated with Cpd A and AZ13581837 compared to vehicle control, albeit not significantly due to the high variability between donors (**Fig. 6E**). The perdurance of the agonist effects following washout may be due to FFAR4 signaling post receptor internalization from intracellular compartments as has been proposed for other GPCRs [56].

**Figure 6.**
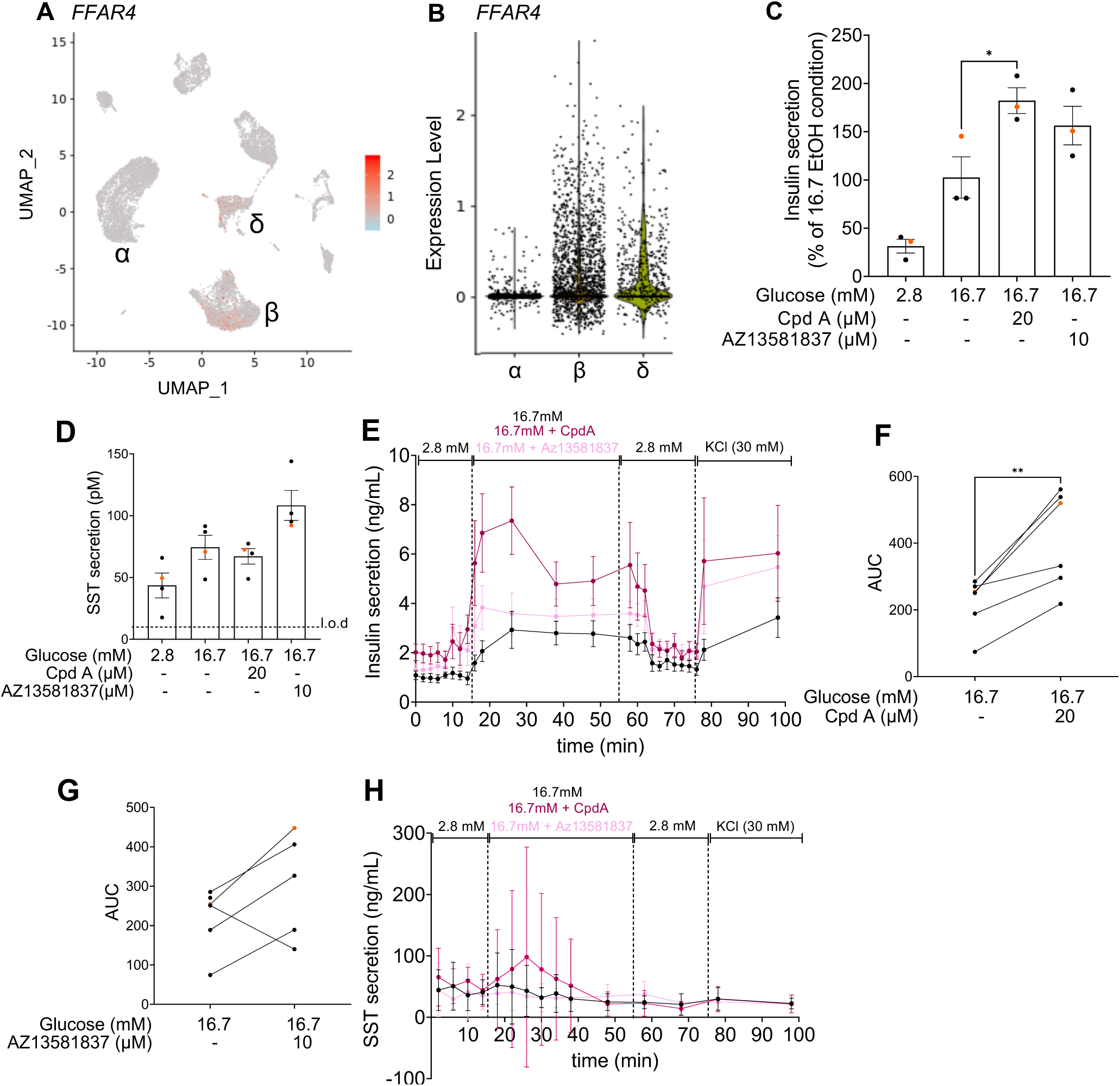
FFAR4 agonists potentiate GSIS but do not inhibit SST secretion in human islets. (A) UMAP plot of five integrated human pancreatic islet single-cell RNA-seq studies as described in [44] showing *FFAR4* expressing cells. (B) Violin plots showing *FFAR4* expression level of individual α, β and δ cells. Adapted from [44]. Insulin secretion presented as a percentage of 16.7-EtOH condition (C) and SST secretion (D) measured in parallel was assessed in 1-h static incubations in response to 2.8 or 16.7 mM glucose with Cpd A (20 μM), AZ13581837 (AZ, 10 μM) or vehicle (EtOH) in human islets. Data are expressed as mean ± SEM from 3-4 donors. Insulin (E) and SST (H) secretion was assessed in human islet perifusions in response to sequential exposure to 2.8 mM glucose, 16.7 mM glucose with or without Cpd A (20 μM) or AZ13581837 (10 μM), followed by 2.8 mM glucose and then 30 mM KCl. Calculation of area under the curve (AUC) from 10 to 50 min for insulin secretion with or without Cpd A (F) or AZ13581837 (G). Data are expressed as mean ± SEM (E, H) or individual values (F, G) from 5-6 donors. ∗p < 0.05, ∗∗p < 0.005 vs 16.7-EtOH condition following one-way ANOVA with Tukey’s post hoc adjustment for multiple comparisons. Male (black); Female (orange). L.o.d, limit of detection.

In line with the static incubation data (**Fig. 6D**), we did not observe an increase in SST secretion in response to glucose and the addition of Cpd A or AZ13581837 was without effect **(Fig. 6H**).

### 3.6 Cpd A stimulates Ca^++^ transients in human β cells

To investigate signaling downstream of FFAR4 in human β cells, human islets were transduced with an adenovirus expressing the Ca^++^ reporter GcaMP6s. Although that GcaMP6s reporter is driven by a ubiquitous promoter (CMV), the reporter signal is predominantly of β cell origin due to their abundance in human islets and the tropism of the viral capsid for β cells [57]. Increasing the glucose concentration from 2.8 to 11 mM stimulated Ca^++^ signaling and subsequent addition of Cpd A led to a further increase (**Fig. 7 –** note that each line in this figure is a whole-islet trace). Following agonist washout, the subsequent addition of the competitive SST receptor antagonist cycloSST increased Ca^++^ fluxes suggesting that SST, albeit not regulated by FFAR4 activation, nevertheless inhibits Ca^++^ signaling in β cells, consistent with a paracrine negative effect on insulin secretion.

**Figure 7.**
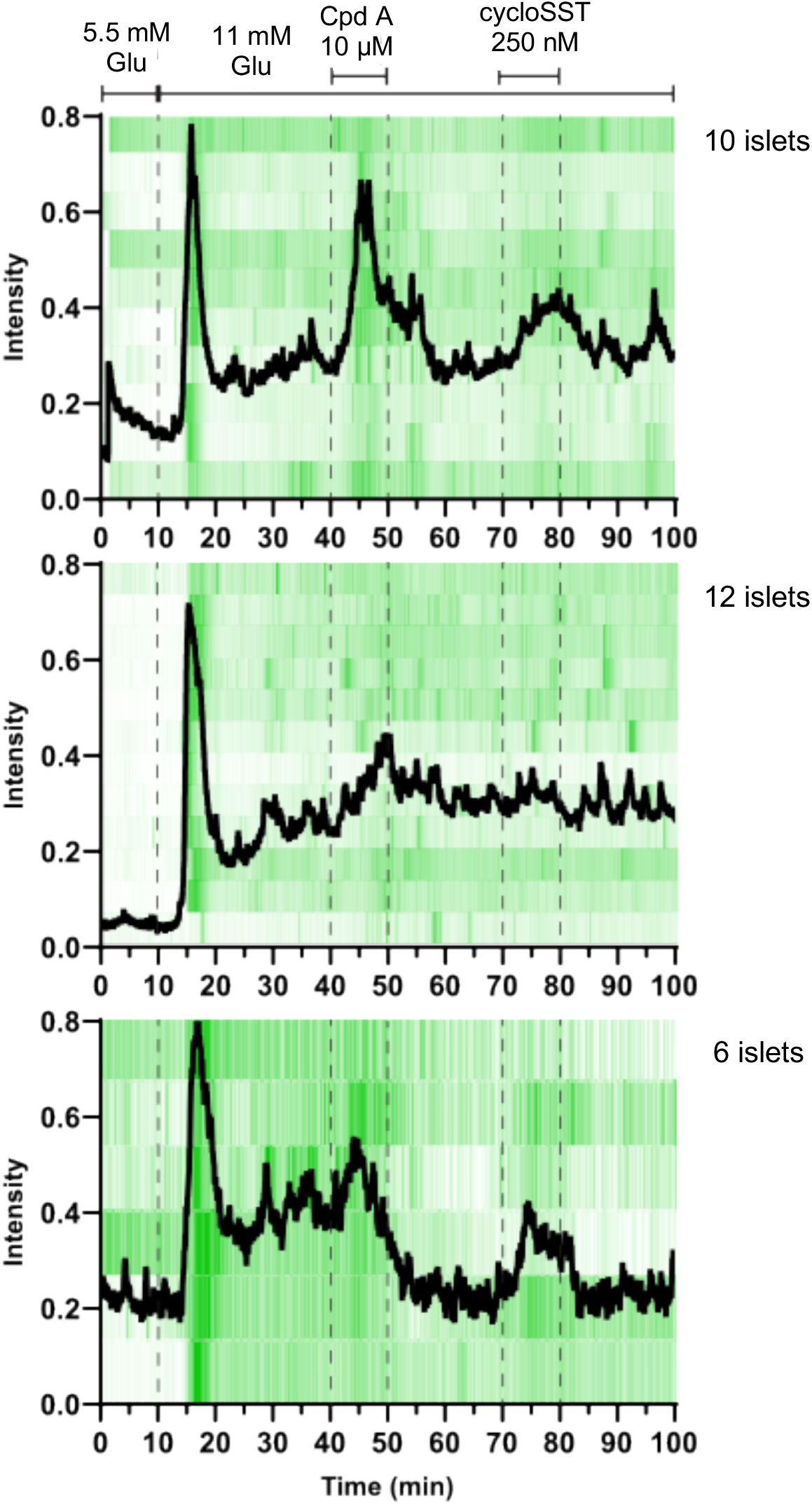
Cpd A stimulates intracellular Ca^++^ transients in human islets. The Ca^++^ sensor GcaMP6s was used to measure Ca^++^ transients in human islets exposed to 5.5 mM glucose followed by 11 mM glucose. Cpd A (10 μM) and cycloSST (250 nM) were added at the times indicated. Panels show Ca^++^ traces for each islet and the average signal for each of 3 technical replicates from one donor (HP-24284-01).

### 3.7 FFAR4 activation potentiates insulin secretion in human EndoC-βH5 cells

To further assess a direct effect of FFAR4 activation on human β cells, we measured insulin secretion in response to FFAR4 agonists in human insulin-secreting EndoC-βH5 cells [47]. Although at the concentration tested the effect of Cpd A did not reach statistical significance, AZ13581837 significantly potentiated GSIS (**Fig. 8**), indicating that FFAR4 agonists act directly on human β cell-like cells to promote insulin secretion.

**Figure 8.**
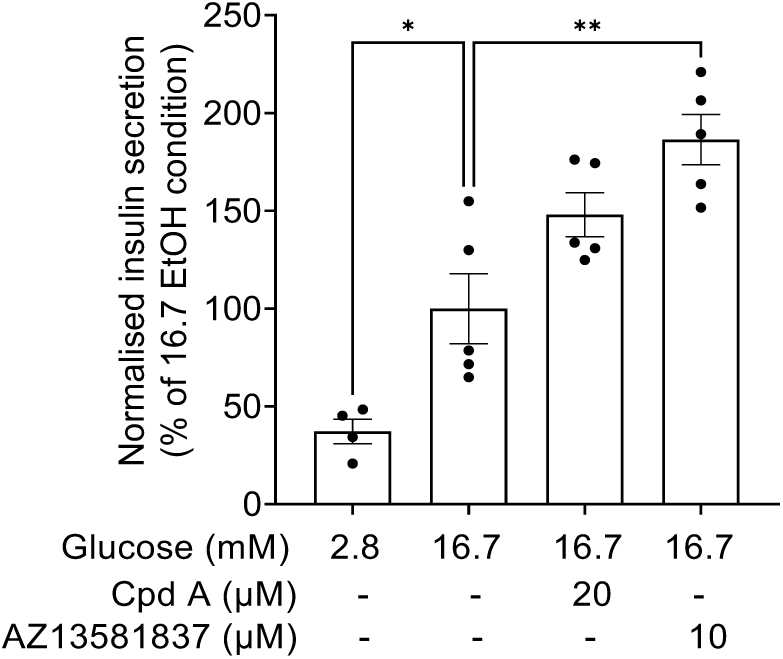
FFAR4 activation potentiates GSIS in human EndoC-βH5 cells. Normalized insulin secretion presented as a percentage of the control 16.7-EtOH condition was assessed in 1-h static incubations in response to 2.8 or 16.7 mM glucose with Cpd A (20 μM), AZ13581837 (10 μM), or vehicle (EtOH) in EndoC-βH5 cells. Data are expressed as mean ± SEM from 4-5 independent experiments. ∗p < 0.05, ∗∗p < 0.005 vs 16.7-EtOH condition following one-way ANOVA with Tukey’s post hoc adjustment for multiple comparisons.

## 4. DISCUSSION

This study was designed to address 3 questions: **1-** What is the relative importance of δ cells and SST in the insulinotropic effect of FFAR4 in mouse islets? **2-** Which G protein does FFAR4 couple to in mouse δ cells? **3-** Does FFAR4 stimulate insulin secretion by similar mechanisms in mouse and human islets? We showed that in the absence of δ cells or SST, FFAR4 agonists no longer potentiate GSIS in mouse islets. In line with an indirect insulinotropic mechanism, FFAR4 agonists have no effect on GSIS in purified mouse β cells. In contrast, FFAR4 agonism directly inhibits SST secretion from purified mouse δ cells, an effect that requires the G protein Gα_z_ as deletion of *Gnaz* abrogates the inhibitory effect on Ca^++^ fluxes and SST secretion. In contrast, in human islets FFAR4 agonists potentiate GSIS without modulating SST secretion, increase Ca^++^ fluxes in β cells, and potentiate GSIS in EndoC-βH5 cells. Overall, our data suggest that the insulinotropic effect of FFAR4 is mediated by distinct mechanisms in mouse and human islets. In mouse islets, by inhibiting SST secretion via FFAR4-Gα_z_ coupling in δ cells, FFAR4 agonists mediate a permissive insulinotropic response. In contrast, in human islets FFAR4 appears to act directly on β cells to potentiate GSIS.

The absence of a potentiating effect of Cpd A on GSIS when δ cells or *Sst* were removed from mouse islets suggests that the effect of FFAR4 agonism is largely indirect. Taken together with our earlier studies [31], these data using complementary models indicate that the insulinotropic effect of Cpd A is mediated by FFAR4-dependent inhibition of δ cell SST secretion. Consistent with an indirect role, Cpd A did not potentiate GSIS in sorted mouse β cells suggesting that FFAR4 signaling is not operative in primary murine β cells. However, *Ffar4* is detected at lower level in murine β cells [31; 32; 35; 36; 39; 40] compared to δ and α cells, and FFAR4 protein was reported to localize to the primary cilium in mouse islet β cells [27]. And in contrast to the lack of effect of Cpd A in purified β cells in our study, Wu et al. [27] and McCloskey et al. [30] demonstrated potentiation of GSIS in murine MIN6 cells in response to the FFAR4 agonists AZ13581837 and TUG891 and in rat BRIN-BD11 insulinoma cells in response to the FFAR4 agonists Cpd A and GSK137647, respectively. This discrepancy could be due to differences between transformed insulin-secreting cells and primary β cells. In addition, given that FFAR4 agonists exhibit ligand-dependent receptor conformational bias resulting in selective G protein coupling, ‘biased signaling’ [58], we cannot exclude that a minor, direct role of FFAR4 in rodent β cells in the control of insulin secretion may be detected in response to other ligands.

Previously we found that pertussis toxin pre-treatment does not prevent Cpd A-mediated repression of GSSS in mouse islets [^31^], suggesting that the inhibitory G proteins Gα_i/o_ are not involved. We therefore considered the alternative possibility that the pertussis toxin-insensitive inhibitory G protein Gα_z_, which is expressed in mouse islets and mediates the inhibition of GSIS by prostaglandin E via the EP3 receptor [59], might be involved. Accordingly, in our previous study we showed that Cpd A reduces forskolin-induced cyclic AMP levels in mouse δ cells [^31^], consistent with the known role of Gα_z_ in the inhibition of adenylate cyclase activity [^59^]. By analyzing *Gnaz^-/-^* mice we showed that Gα_z_ mediates FFAR4-dependent inhibition of Ca^++^ signaling and SST secretion in δ cells. In contrast, Stone et al. [36] provided evidence that repression of GSSS by the FFAR4 agonist Metabolex 36 is pertussis toxin-sensitive in isolated mouse islets. Both Gα_z_ and Gα_i/o_ are expressed in δ cells [^39^], and mouse FFAR4 has been shown previously to couple to Gα_i/o_ [^32^]. These discrepant results could again be explained by biased signaling [^58^], whereby Cpd A favours FFAR4-Gα_z_ coupling whereas Metabolex 36 favours FFAR4-Gα_i/o_ coupling in mouse δ cells. On the other hand, in heterologous cells Cpd A promotes FFAR4-Gα_q_ coupling [^5^] and in BRIN-BD11 cells Cpd A promotes GSIS [^30^], suggesting possible, context-dependent coupling with stimulatory G proteins (eg. Gα_q_ or Gα_s_).

Why Cpd A favours inhibitory FFAR4-Gα_z_ coupling in δ cells where Gα_q_ is also expressed [39] may be due to cell-type-specific factors, other than differential G protein expression, including interference from other δ-cell GPCRs that modulate the availability of G protein isoforms at FFAR4, as suggested from GPCR competition studies [60].

Unexpectedly, we found that Cpd A still potentiates GSIS despite the absence of Cpd A-mediated repression of SST secretion in *Gnaz^-/-^* mouse islets. These data could be explained by competitive coupling of FFAR4 to Gα_z_ or Gα_q_ in β cells whereby in wild-type β cells neither inhibitory Gα_z_ or stimulatory Gα_q_ signaling predominates in response to Cpd A leading to a net null effect on insulin secretion, whereas in *Gnaz^-/-^* β cells Cpd A signals exclusively via FFAR4-Gα_q_ coupling to potentiate GSIS. Further studies are required to test this hypothesis.

In human islets, we found that FFAR4 agonists stimulate Ca^++^ fluxes and insulin secretion in the absence of detectable, agonist-regulated SST secretion. Accordingly, the FFAR4 agonists AZ13581827 and TUG891 potentiate GSIS in human islets and pseudoislets in static incubations [27]. Although to our knowledge, ours is the first study addressing a role of FFAR4 agonists in SST secretion in human islets, Du et al. [32] found that the FFAR4 inverse agonist AH7614 increases GSSS in human islets, suggesting that FFAR4 signaling, possibly due to endogenous islet-derived FFAR4 ligands, inhibits SST secretion. In light of these data, and given the limitations of the static and perifusion assay to accurately detect small changes in SST levels, we cannot exclude a minor contribution of FFAR4-dependent regulation of SST secretion in the insulinotropic response to FFAR4 agonists in human islets. Nevertheless, *FFAR4* expression is elevated in both human β and δ cells [41; 43; 44], suggesting a greater contribution of FFAR4 in human β cells compared to mouse where expression in β cells is relatively weak compared to δ cells [31; 32; 35; 36; 39-41]. In support of a potential direct role of FFAR4 in human β cells, here we showed that FFAR4 agonists potentiate GSIS in EndoC-βH5 cells and stimulates Ca^++^ in human β cells. Although FFAR4 exhibits promiscuous coupling with Gα_i/o_, Gα_s_ and Gα_q_ and β-arrestin in heterologous cells [^42^], the direct effect of Cpd A on GSIS suggests a coupling to stimulatory G proteins Gα_s_ and/or Gα_q_ in this context. Further studies will be required to identify the coupling of FFAR4 in human β cells.

In conclusion, our data suggest that FFAR4 controls insulin secretion by distinct regulatory mechanisms in rodent and human islets. FFAR4 activation permits insulin secretion from mouse islets indirectly via Gα_z_-coupled inhibition of δ−cell SST secretion, while in human islets, it stimulates insulin release via a direct effect on β cells. These key species-related differences are to be taken into account as FFAR4 is considered a potential therapeutic target for metabolic diseases.

## ACKNOWLEDGEMENTS

This study was supported by the U.S. National Institutes of Health (R01-DK-132597 to M.O.H. and V.P.) and a Discovery Grant from the Natural Sciences and Engineering Research Council of Canada (RGPIN-2016-03952 to V.P.). L.R. was supported by a fellowship from the Société Francophone du Diabète. Human pancreatic islets were provided by the NIDDK-funded Integrated Islet Distribution Program (IIDP) (RRID:SCR _014387) at City of Hope, NIH Grant # U24DK098085. The Graphical Abstract was created with BioRender.com.

We thank Maria Sörhede Winzell and Linda Sundström from Astra Zeneca for providing AZ13581837; Mélanie Éthier, Sabrina Sedano-Benitez, and Gaël Dulude from the CRCHUM for valuable technical assistance; Ryan Hart from Mark Huising’s laboratory at UC Davis for help with Ca^++^ imaging and Raphaël Scharfmann for help with the islet cell sorting protocol.

## 5. AUTHOR CONTRIBUTIONS

**Laura Reininger**: Conceptualization, methodology, investigation, formal analysis, and writing original draft. **Muhammad Rehman**: Methodology, investigation, formal analysis, and visualization. **Amélia Bouabcha**: Investigation. **Sarah Ferragne**: Investigation. **Caroline Tremblay**: Methodology and investigation. **Mélanie Ethier**: Methodology. **Julien Ghislain**: Conceptualization, validation, writing, review, editing, supervision, and project administration. **Michelle E. Kimple**: Methodology and editing. **Mark O. Huising**: Conceptualization, validation, writing, review, editing, supervision, funding acquisition, and project administration. **Vincent Poitout**: Conceptualization, validation, writing, review, editing, supervision, funding acquisition, and project administration.

## GLOSSARY

Ad: Adenovirus
AMP: Adenosine monophosphate
AUC: Area under the curve
Cpd A: Compound A
CMV: Cytomegalovirus
DT: Diphtheria toxin
DTR: Diphtheria toxin receptor
FC: Fold change
FFAR4: Free fatty acid receptor 4
GPCR: G protein-coupled receptor
GSIS: Glucose-stimulated insulin secretion
GSSS: Glucose-stimulated somatostatin secretion
SST: Somatostatin

