## Supplementary Material for "Free fatty acid receptor 4 agonists stimulate insulin secretion via different mechanisms in mouse versus human islets"

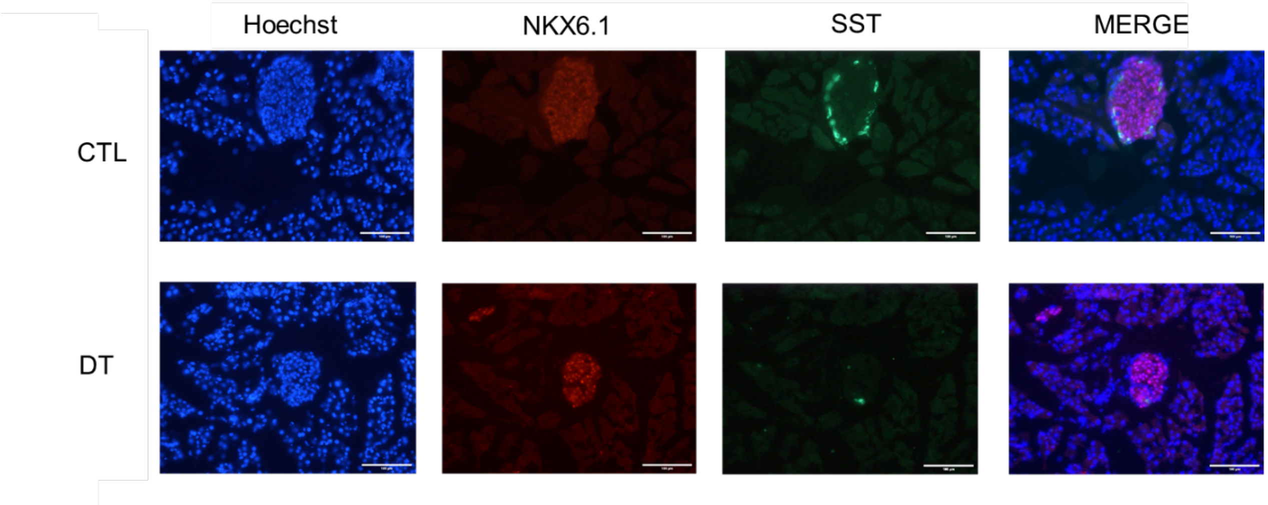


**Supplementary Figure S1. Immunofluorescent staining of pancreatic sections from control and δ-cell-deficient male mice.** Pancreatic section from saline (CTL) or DT injected *Sst*$^{+/cre}$;*ROSA26*$^{+/idtr}$ mice (DT) were stained for NKX6.1 (red), SST (green) and nuclei (blue). Representative images from n=3 are shown. Scale bar, 100 µm.

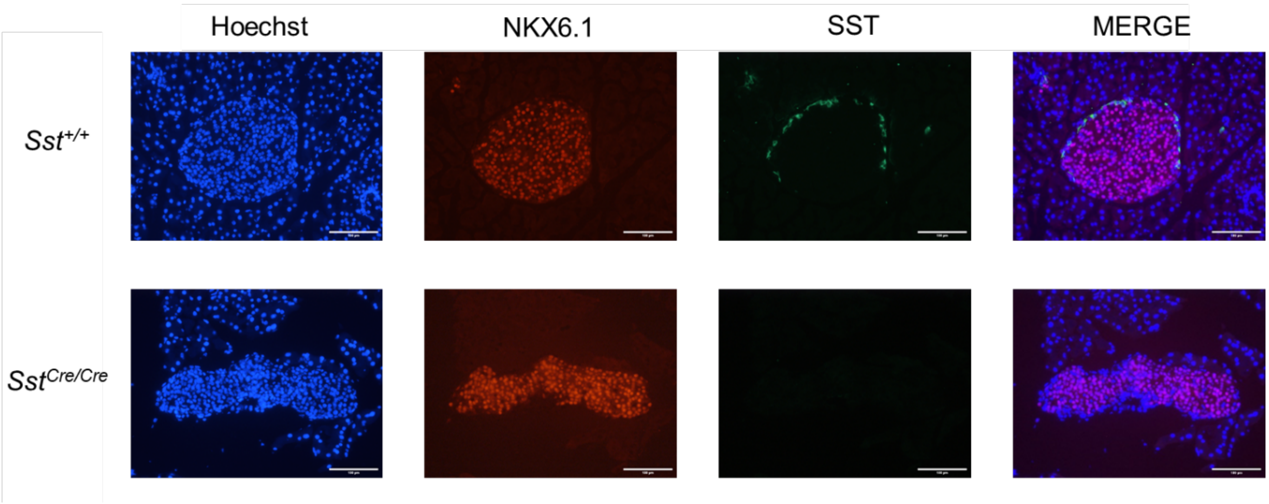


**Supplementary Figure S2. Immunofluorescent staining of pancreatic sections from control and SST-deficient male mice.** Pancreatic section from *Sst*$^{+/+}$ and *Sst*$^{cre/cre}$ mice were stained for NKX6.1 (red), SST (green) and nuclei (blue). Representative images from n=3 are shown. Scale bar, 100 µm.

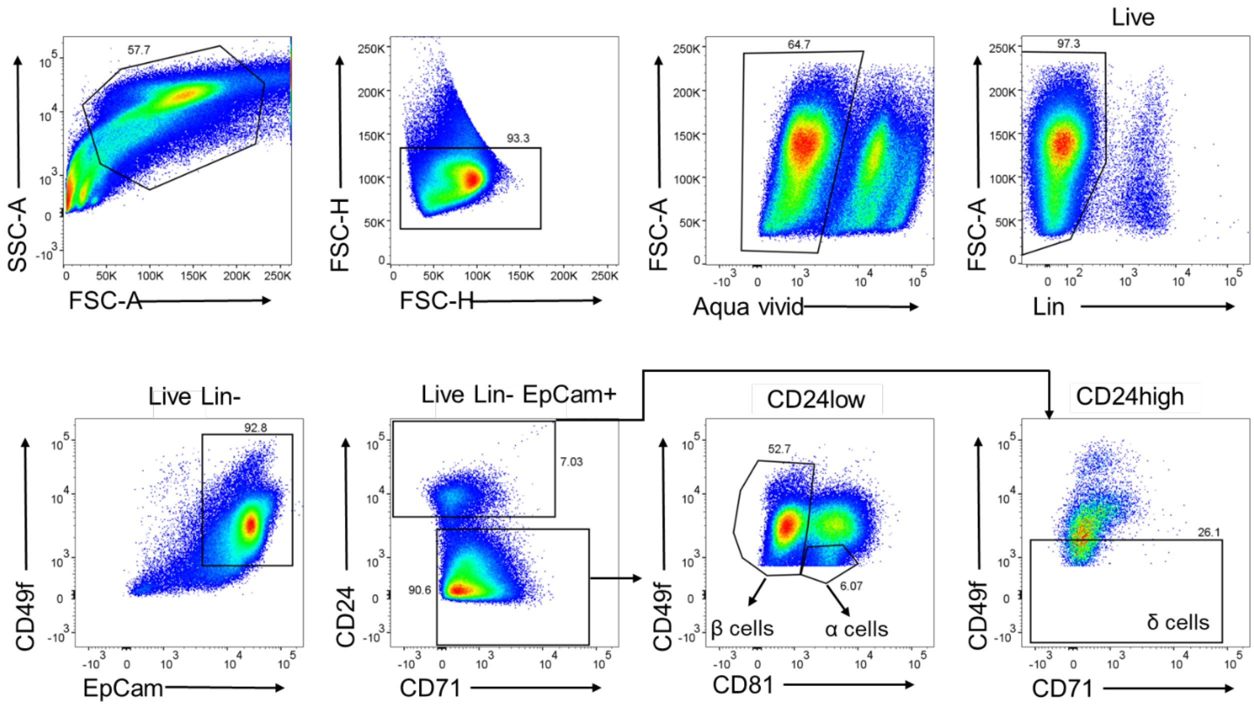


**Supplementary Figure S3. Flow cytometric sorting of α, β and δ cells from male mouse islets.** Representative plots of dispersed wild-type male mouse islets. Surface markers CD24, CD71, CD49f and CD81 expression were analyzed on $EpCam^{+}$ cells after excluding doublets, dead cells (Aqua Vivid) and endothelial cells ($Lin^{-}$: $CD45^{-}$, $TER119^{-}$ and $CD31^{-}$). $EpCam^{+}$ cells were divided into $CD24^{low}$ and $CD24^{high}$ cell fractions. $CD24^{low}$ cells were further subdivided into $CD81^{-}$ (β cell fraction) and $CD81^{+}/CD49f^{-}$ (α cell fraction). Among $CD24^{high}$ cells, $CD49f^{-}$ cells were selected (δ cell fraction).

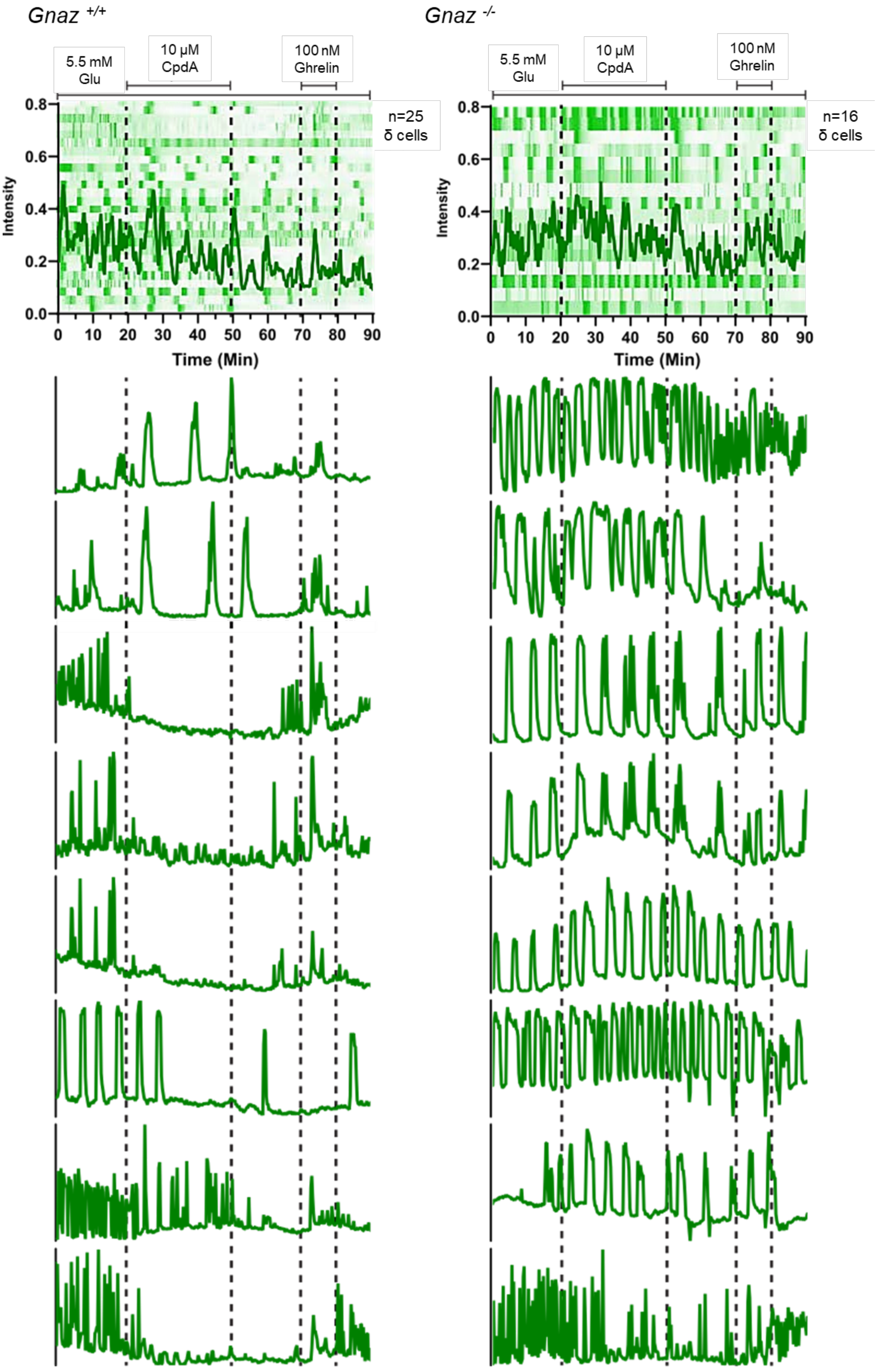

Gnaz +/+
Gnaz -/-
5.5 mM Glu
10 µM CpdA
100 nM Ghrelin
n=25 δ cells
n=16 δ cells
Intensity
Time (Min)

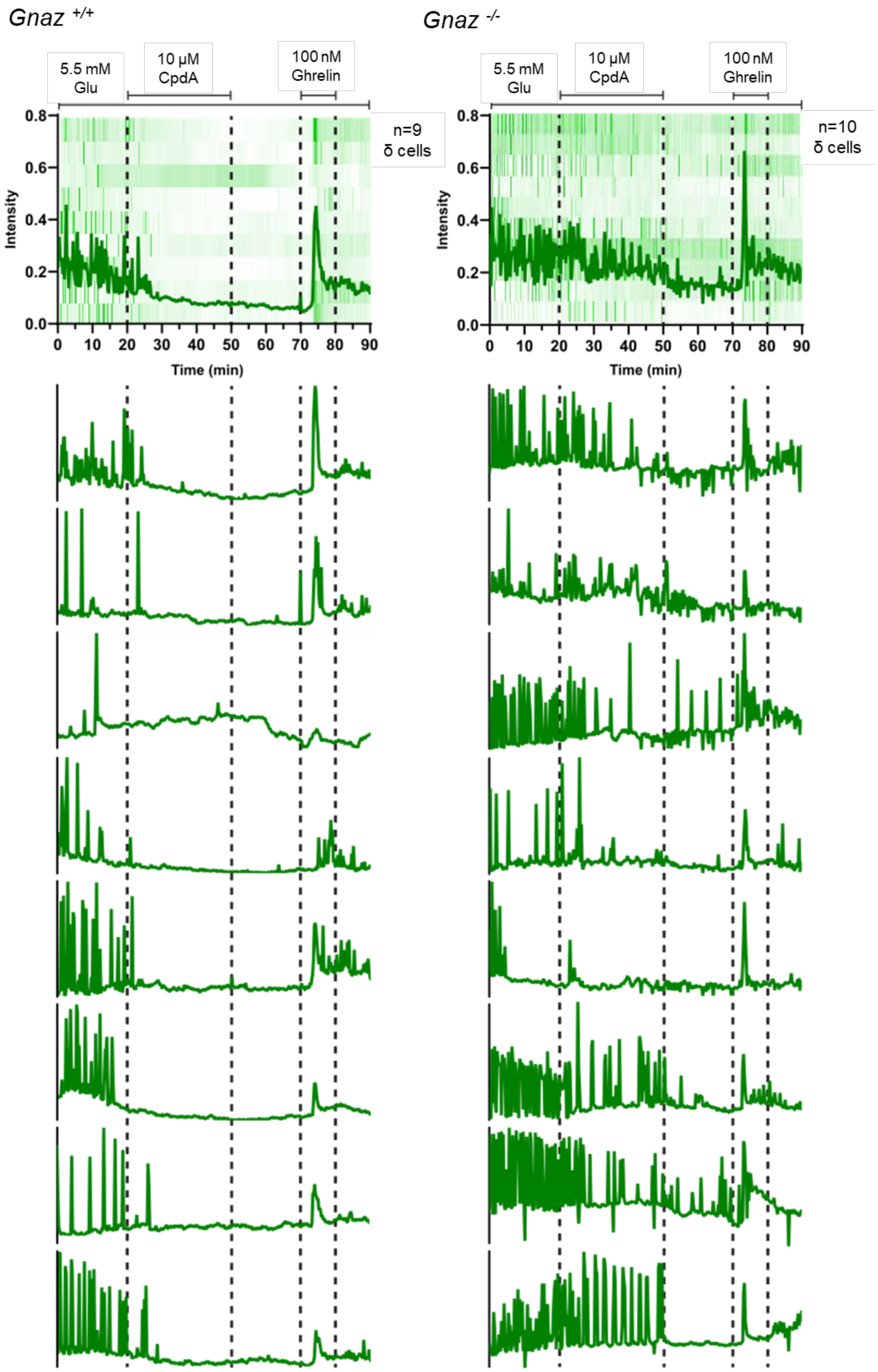

Gnaz +/+
Gnaz -/-
5.5 mM Glu
10 µM CpdA
100 nM Ghrelin
n=9 δ cells
n=10 δ cells
Intensity
Time (min)

**Supplementary Figure S4.** ***Gnaz*** **deletion abrogates CpdA-mediated decrease in intracellular** $\mathbf{Ca^{++}}$ **transients in male mouse δ cells.** The $Ca^{++}$ sensor jRGECO1b was used to measure $Ca^{++}$ transients in individual δ cells in the presence of 5.5 mM glucose in $Gnaz^{+/+}$ (left) and $Gnaz^{-/-}$ (right) male mouse islets. Cpd A (10 μM) was added at the times indicated. Ghrelin (100 nM)-induced $Ca^{++}$ signaling served as a positive control to identify δ cells. Aggregate data of all δ cells analyzed plotted with the average intensity (upper panels) and traces of individual δ cells (lower panels) are shown. Data represent the results of the analysis of 9-25 δ cells in each of 2 batches of $Gnaz^{+/+}$ and $Gnaz^{-/-}$ islets.

**SUPPLEMENTARY TABLES**

**Supplementary Table 1: Primers used for genotyping.**

| Gene | Primer 1 | Primer 2 | Primer3 |
|---|---|---|---|
| *Gnaz*$^{-/+}$ | CTCTCTCATCTGCCCGTCACAC | TACTCCTTGCAGGCGTCCAGGTT | GGTGGATGTGGAATGTGTGCGA |
| *Sst*$^{+/Cre}$ | GGGCCAGGAGTTAAGGAAGA | TCTGAAAGACTTGCGTTTGG | TGGTTTGTCCAAACTCATCAA |
| *ROSA26*$^{+/\ iDTR}$ | AAAGTCGCTCTGAGTTGTTAT | GGAGCGGGAGAAATGGATATG | GCGAAGAGTTTGTCCTCAACC |

**Supplementary Table 2: Sources of antibodies used for flow cytometry.**

| Fluorophore | Antibody | Dilution | Reference |
|---|---|---|---|
| PerCP/Cyanine5.5 | Anti-mouse TER-119 | 1/100 | BioLegend; 116228 |
| PerCP/Cyanine5.5 | Anti-mouse CD31 | 1/100 | BioLegend; 102522 |
| PerCP/Cyanine5.5 | Anti-mouse CD45 | 1/100 | BioLegend; 103132 |
| APC/Fire™ 750 | Anti-mouse CD24 | 1/100 | BioLegend; 101840 |
| BV605 | Anti-mouse CD326 (Ep-CAM) | 1/100 | Biolegend; 118227 |
| PE-Cyanine7 | Anti-mouse CD71 | 1/100 | BioLegend; 113812 |
| PE | Anti-human/mouse CD49f | 1/200 | BioLegend; 313612 |
| BV421 | Anti-mouse/rat CD81 | 1/100 | BD biosciences; 740060 |

**Supplementary Table 3: Primers used for qPCR.**

| RNA | Forward | Reverse |
|---|---|---|
| Ins1 | CAGAGACCATCAGCAAGCAG | GGGACCACAAAGATGCTGTT |
| Gcg | TGTCTACACCTGTTCGCAGC | AATCCAGGTGTTGTGACTGC |
| Sst | GCTGAAGGAGACGCTACCGA | GGGGGCCAGGAGTTAAGGAA |
| Ppia | TCTGAGCACTGGAGAGAAAGG | TTCTCTCCGTAGATGGACCTG |

**Supplementary Table 4: Sources of antibodies used for immunohistochemistry.**

| Antibody | Dilution | Reference |
|---|---|---|
| Anti-mouse NKX6-1 | 10 µg/ml | DSHB; F55A12 |
| Anti-mouse SST | 1/200 | Abcam; ab108456 |
| Cy3 Goat anti-mouse | 1/400 | Thermo Fisher Scientific; A10521 |
| Alexa Fluor 488 Donkey anti-rabbit | 1/500 | Thermo Fisher Scientific; A21206 |

**Supplementary Table 5: Human islet donor information.**

**ADI, Alberta Diabetes Institute; IIDP, Integrated Islet Distribution Program; SI, stimulation index**

| | **1** | **2** | **3** | **4** | **5** | **6** | **7** | **8** | **9** | **10** |
|---|---|---|---|---|---|---|---|---|---|---|
| **Unique identifier** | SAMN40435627 | SAMN25860453 | SAMN23958504 | R421 | HP-24122-01 | HP-24203-01 | HP-24254-01 | HP-24265-01 | HP-24269-01 | HP-24284-01 |
| **Age (years)** | 32 | 44 | 42 | 60 | 38 | 65 | 57 | 43 | 44 | 55 |
| **Sex** | M | M | M | F | M | M | M | M | M | F |
| **BMI (kg/m$^2$)** | 41.7 | 28.5 | 37.9 | 25.9 | 29.2 | 33 | 28.7 | 27.1 | 27.2 | 23.3 |
| **HbA1c** | 5.3 | 6 | 5.7 | 5.4 | 5.2 | 5.4 | 5.9 | 5.3 | 5.5 | 5.5 |
| **Origin** | IIDP | IIDP | IIDP | ADI IsletCore | Prodo Labs. | Prodo Labs. | Prodo Labs. | Prodo Labs. | Prodo Labs. | Prodo Labs. |
| **Islet isolation center** | Southern California Islet Cell Resource Center | UPenn Islet Transplant Center | Prodo Labs. | ADI IsletCore | Prodo Labs. | Prodo Labs. | Prodo Labs. | Prodo Labs. | Prodo Labs. | Prodo Labs. |
| **Diabetes** | No | No | No | No | No | No | No | No | No | No |
| **Cause of death** | Head trauma | Head trauma | Stroke | Not reported | Anoxic event | Stroke | Stroke | Head trauma | Head trauma | Stroke |
| **Warm ischemia time** | Not reported | Not reported | Not reported | Not reported | Not reported | Not reported | Not reported | Not reported | Not reported | Not reported |
| **Cold ischemia time** | 6h59 min | 7h58 min | 9h54 min | 18h | Not reported | Not reported | Not reported | Not reported | Not reported | Not reported |
| **Estimated purity (%)** | 80 | 90 | 90 | 95 | 95 | 90 | 95 | 90 | 95 | 85 |
| **Estimated viability (%)** | 95 | 93 | 95 | 73 | 95 | 95 | 95 | 95 | 95 | 95 |
| **GSIS (SI)** | 1.5 | 1.5 | 1.8 | 1.7 | 3.1 | 1.1 | 3.1 | 2.6 | 1.1 | 1.3 |
